# Auxin Response Factors — output control in auxin biology

**DOI:** 10.1101/129122

**Authors:** Mark Roosjen, Sébastien Paque, Dolf Weijers

**Affiliations:** Laboratory of Biochemistry, Wageningen University, Stippeneng 4, 6708WE Wageningen, the Netherlands

## Abstract

The phytohormone auxin is involved in almost all developmental processes in land plants. Most, if not all, of these processes are mediated by changes in gene expression. Auxin acts on gene expression through a short nuclear pathway that converges upon the activation of a family of DNA-binding transcription factors. These AUXIN RESPONSE FACTORS (ARFs) are thus the effector of auxin response and translate the chemical signal to the regulation of a defined set of genes. Given the limited number of dedicated components in auxin signaling, distinct properties among the ARF family likely contributes to the establishment of multiple unique auxin responses in plant development. In the two decades following the identification of the first ARF in *Arabidopsis* much has been learnt about how these transcription factors act, and how they generate unique auxin responses. Progress in genetics, biochemistry, genomics and structural biology have helped to develop mechanistic models for ARF action. However, despite intensive efforts, many central questions are yet to be addressed. In this review we highlight what has been learnt about ARF transcription factors, and identify outstanding questions and challenges for the near future.

---

In the past decade, the auxin signaling pathway that leads to gene expression responses has been characterized in detail (Weijers and Wagner, 2016). The core of the auxin pathway, which takes place in the nucleus, is centered around three different factors (Figure 1). The pathway relies on the inhibiting role of Aux/IAAs, inhibitors of the Auxin Response transcription Factors (ARFs) that allow auxin-dependent gene expression. To unlock the system, auxin binds directly to the SCF (TIR1/AFB) ubiquitin ligase and hence increases the affinity for Aux/IAAs proteins, leading to their subsequent degradation by the 26S proteasome. Released from Aux/IAA inhibition, ARFs can then modulate auxin-dependent gene transcription. Based on this model, ARFs are considered as the output of the nuclear auxin pathway.

**Figure 1:**
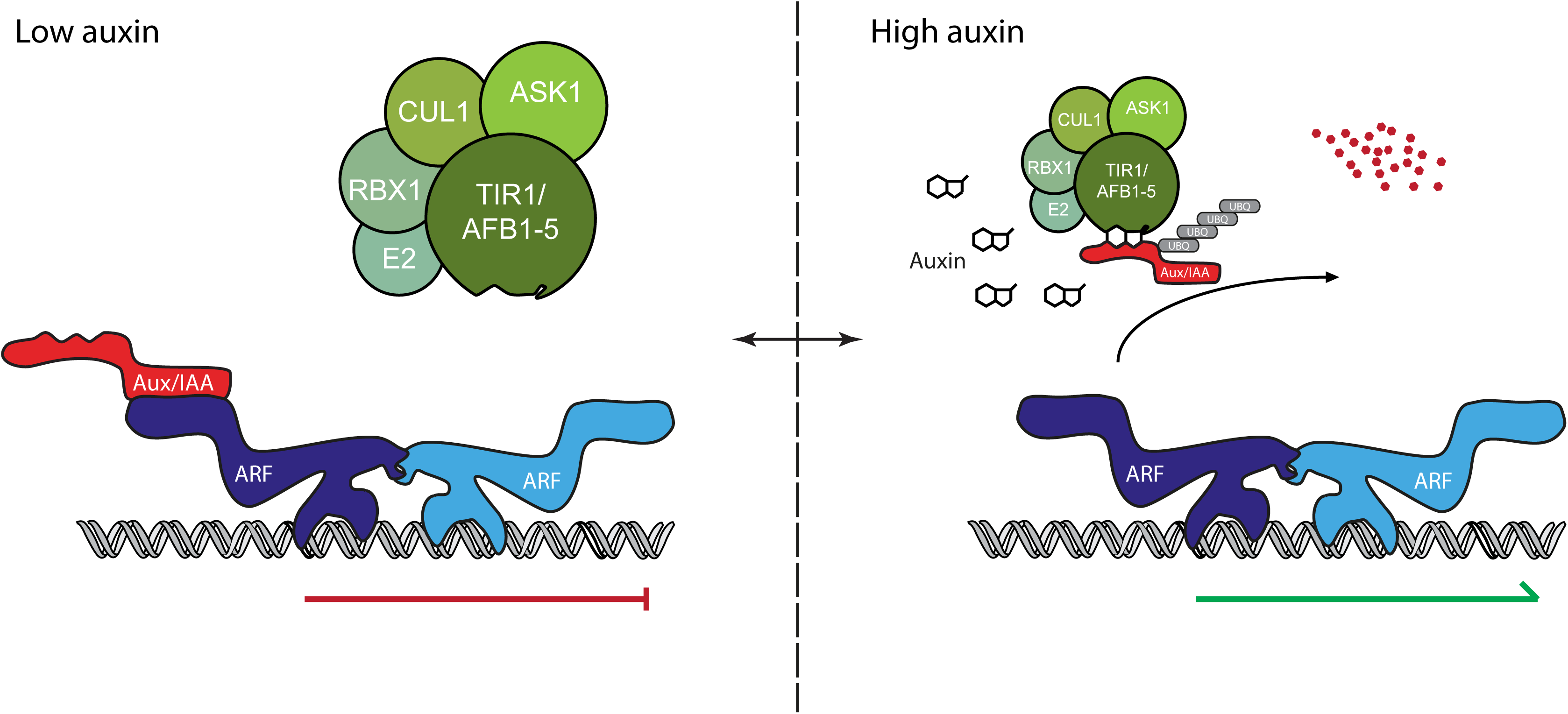
The nuclear auxin pathway. Regulation of auxin output is executed by ARFs. Under low auxin levels, the Aux/IAA transcriptional co-repressors prevent ARFs from controlling auxin-regulated genes. When auxin levels increase, auxin serves as “molecular glue” between the TIR1/AFB receptor and the Aux/IAA protein. This leads to subsequent ubiquitination and degradation of the Aux/IAAs, releasing ARFs from inhibition. *Protein abbreviations: ARF, AUXIN RESPONSE FACTOR; ASK1, ARABIDOPSIS SKP1 HOMOLOGUE; Aux/IAA, AUXIN/INDOLE-3-ACETIC ACID; CUL1, CULLIN 1; RBX1, RING-BOX 1; TIR1/AFB, TRANSPORT INHIBITOR RESISTANT 1/AUXIN SIGNALING F-BOX.*

To date, these three signaling components appear to be sufficient to trigger nuclear auxin signaling in a heterologous system (Pierre-Jerome *et al.*, 2014). The fact that these three components belong to multigene families offers some explanations for how such a simple pathway can control such a wide array of different developmental processes. Importantly, there may be significant functional specialization among ARFs. However, the precise mechanisms that generate dynamics and specificity to auxin output are largely unknown, but the community is currently addressing this challenge. This review will focus on the effectors of the nuclear auxin pathway in *Arabidopsis*. Given their position in the auxin pathway, we focus our discussion on the mode of action of the ARFs. Recent insights in the past years have allowed the community to see these transcription factors in a new light. This review will give a comprehensive overview of the work that has been done and will raise questions that need to be tackled in the future.

## Domain organization of ARF transcription factors

The *Arabidopsis* genome encodes 23 ARFs that fall into three subclasses called A, B and C. Importantly, only few loss of function mutants show an obvious growth phenotype, and double mutants have revealed gene redundancy between close relatives (Okushima *et al.*, 2005). However, a combination of promotor-swap, misexpression and loss-of-function approaches suggested that ARFs are not interchangeable and lead to specific phenotypes (Rademacher *et al.*, 2011, 2012). Most ARFs share a similar topology with three conserved protein domains and the properties of these need to be understood in detail. Here, the three representative domains will be introduced separately.

All ARFs possess at their N terminus a conserved DNA binding domain (DBD) (Okushima *et al.*, 2005; Boer *et al.*, 2014). Surprisingly, a phylogenetic tree using only DBD protein sequences appears similar to that using full-length protein sequences (Boer *et al.*, 2014). This suggests that some functional specificities could be provided by this domain. Crystal structures of the DBDs of ARF1 and ARF5 revealed an unique 3D conformation of the DBD and highlight the presence of three different subdomains: a B3 subdomain showing similarity with the DNA-contacting domain of bacterial endonucleases, a dimerization domain (DD) allowing ARF dimerization and a Tudor like ancillary domain (AD) of unknown function which might be involved in an interaction with the DD. The DBD of ARFs fulfils a critical role for a transcription factor: Recognition of a DNA motif, called the auxin responsive element (AuxRE). In addition, the DBD allows dimerization of ARFs that mediates biological activity.

### Specific DNA binding through the DNA-binding domain

One of the functions of a transcription factor is to bind DNA with sequence specificity. The B3 subdomain is involved in the recognition of the ARF-specific AuxRE DNA motif. The crystal structures of the DBD of ARF1 and ARF5 homodimers, as well as the complex of ARF1 DBD with DNA allowed to visualize the mode of protein-DNA interaction. This ARF-DNA crystal confirmed results obtained two decades ago when domains involved in ARF DNA binding had been discovered (Ulmasov *et al.*, 1997a) and shows how amino acids in the DBD interact with the DNA binding motif TGTCTC (Boer *et al.*, 2014). Mutations in these DNA-interacting amino acids indeed affect their DNA binding properties and their biological activity.

The canonical TGTCTC was originally identified in promotors of auxin-responsive genes in pea and soybean, and was shown to mediate ARF-activated gene expression (Ulmasov *et al.*, 1995, 1997*a,* 1999*a*). In the past few years, different techniques have broadened the spectrum of known AuxREs. For example, protein-binding microarrays (PBMs) showed that the original AuxRE was not the sequence with the highest ARF-binding affinity, and instead identified the TGTCGG element as a high-affinity binding site (Boer *et al.*, 2014; Franco-Zorrilla *et al.*, 2014). Likewise, TGTCGG also appeared as a representative DNA binding motif of ARF2 and ARF5 in a “cistrome” analysis that measured in vitro binding to genomic fragments (O’Malley *et al.*, 2016).

This higher affinity for TGTCGG has been translated into an optimized artificial auxin response reporter where the 9 TGTCTC repeats in the widespread “DR5” tool have been replaced by TGTCGG repeats (DR5v2) (Liao *et al.*, 2015). This subtle change leads to improvement of the sensitivity of the marker. The coexistence of these two AuxRE’s does not conflict with the numerous results showing the involvement of TGTCTC, but rather enlarge the scope of *cis* elements in auxin response. In fact, the TGTCGG motif appeared to be only present in a third of the strong cistrome peaks of ARF2 and ARF5 and its presence was distinct from the AuxRE sequence TGTCTC (Boer *et al.*, 2014). The significance of AuxRE diversification is still unknown but gene ontology enrichment analysis of genes from auxin transcriptomes suggest that there is a correlation between particular AuxRE’s and specific processes (Zemlyanskaya *et al.*, 2016).

PBMs on ARF1 and ARF5 DBD’s tested all the variants possible from TGTCNN and show that ARFs are in fact able to bind various variants. At the same time, an indirect proof that other TGTCNN variants could be involved in auxin response came from a meta-analysis of auxin transcriptomes published previously (Zemlyanskaya *et al.*, 2016), as well as from cell type specific root transcriptomes (Bargmann *et al.*, 2013). Correlation with auxin up/down regulation and overrepresentation of AuxRE’s highlights putative new AuxRE that will need to be biologically tested. Most of the examples of biological relevance used, as a proof of concept, the canonical AuxRE TGTCTC. e.g. (Weiste and Dröge-Laser, 2014; Ripoll *et al.*, 2015). Understanding the code hidden behind the disposition of AuxREs along the genome is of great importance to understand ARFs mode of action and how auxin responsiveness is specified.

As the crystals structures of ARF1 and ARF5 DBDs show a high degree of similarity, Boer et al. tested the ability of the ARF1 and ARF5 dimers to bind differently spaced AuxREs. Surprisingly, ARF1 and ARF5 did not behave the same regarding the difference in space between two palindromic AuxRE’s. ARF5 seemed to be more lenient than ARF1. This result gave birth to the caliper model where different ARFs can bind different AuxRE motifs with affinity depending on spacer length. This model is supported by the analysis of the cistrome of ARF5 and ARF2 where analysis of the enrichment of AuxRE in promotor of genes bound by the two ARFs show distinct patterns (O’Malley *et al.*, 2016). This caliper theory emphasizes the cooperative binding of two AuxREs where this interaction enhances the binding of the homodimers to DNA compared to binding on the DNA independently (Boer *et al.*, 2014).

In addition to sequences of the AuxRE and the spacing between two AuxRE’s, the orientation of the elements is also an important parameter for binding specificity. Since the discovery of the AuxRE, it is known that differently oriented AuxREs are auxin inducible (Guilfoyle *et al.*, 1998). Cistromes for ARF2 and ARF5 clearly show that both proteins do not bind the same motif (O’Malley *et al.*, 2016). The difference in orientation between direct repeats and inverted repeats should impact the interactions between two AuxREs. The fact that ARF2 and ARF5 do not have the same motifs preferences could reflect specific conformation for homo/hetero dimerization of the ARF on composite AuxREs. However, structural information is at present only available for binding of the ARF1 DBD to an inverted repeat (Boer *et al.*, 2014), and it remains an open question whether alternative dimerization modes underly binding to alternative repeats.

Some correlation seems to exist between the number of AuxRE in a promotor region and its auxin inducibility (Berendzen *et al.*, 2012; O’Malley *et al.*, 2016). If several variants of AuxRE’s confer auxin responsiveness, and the spacing or orientation of AuxRE modules lead to different affinities for the ARFs, it can explain the functional diversity of ARFs and how every ARF could be involved in different developmental processes and why they have specific transcriptomes.

Crystallography of the DBD of ARF1 and ARF5 show that they homodimerize through their DD mediated by hydrophobic interactions. A critical question is whether this homodimerization is biologically relevant. One of the arguments could be that point mutations on amino acids involved in the homodimerization of ARF5 failed to rescue the strong phenotype of the loss of function mutant of ARF5 and without causing any change in the protein folding (Boer *et al.*, 2014). Another piece of evidence to support the biological role of the ARF dimerization is provided by a study in the crop *Brassica napus* where a variant lacking 55 amino acids in the N-terminal domain of ARF18 was unable to dimerize. This dimerization seems to be a requirement for activity, as truncated ARF18 was not able to either bind the DR5 element or inhibit the expression of an auxin response reporter like the wild-type protein (Liu *et al.*, 2015). Moreover, this deletion leads to decreased fruit size and seed weight. While some studies show some heterodimerization between different ARFs, currently it is not known whether the DBD is involved in this interaction.

### Modulating gene activity through the middle region

While the ARF DBD is highly similar in structure and sequence, the middle Region (MR) shows a strongly contrasting property in that it displays the highest divergence in amino acid composition of the ARFs. Thus far, research has primarily focused on the functional properties of the DBD and the PB1 domain, and the properties of the MR have largely remained elusive. However, the MR has offered a framework to categorize the ARF family into either activators or repressors. This classification has been based on the enrichment of specific amino acids in the MR, as well as on the ability of some tested ARFs to either activate or repress transcription from promotors containing the canonical AuxRE TGTCTC (Ulmasov *et al.*, 1999b; Tiwari *et al.*, 2003). The activator/repressor categorization correlates with the division in subgroups A/B/C. Those ARFs tested as activators belong to class A, while class B ARFs encompass the ones tested as repressors (Tiwari *et al.*, 2003).

The class A ARFs, regarded as activators, carries MR’s that are enriched in glutamines, while MR’s in class B and C ARFs have a strong enrichment in serines, prolines and threonines. This observation has not yet gone beyond a correlation, and it is unclear what mechanisms underlie activation and repression. Transient expression experiments of class B ARFs on a few known auxin-dependent promotors did not show a strong gene induction after auxin treatment. However, no genome-wide analysis of transcriptomes has been conducted on class B/C ARFs. It is worth to point out that the promotors used in transient expression assays mainly contained TGTCTC motifs and that, based on the recent knowledge on ARF binding sites preferences, other motifs would perhaps be better suited for analyzing class B/C ARF activity. This should be thoroughly studied to gain better insight into the mode of action of the different classes of ARFs. The important fundamental question of how ARFs function cannot be answered only with a study in heterologous systems on a small set of specific genes. Particularly because genetic studies show that class B and C ARFs can be linked to auxin regulated processes, and that class A ARFs are able to repress certain genes (Sessions and Zambryski, 1995; Sessions *et al.*, 1997; Nemhauser *et al.*, 2000; Zhao *et al.*, 2010; Zhang *et al.*, 2014), the categorization of ARFs into activator and repressor categories should be exercised with caution.

An emerging concept in eukaryotic transcription factor biology is the usage of intrinsic disorder (ID) to elicit specific and rapid conformational changes to allow for adaptive interaction surfaces, conditional DNA binding or modulation of protein function through posttranslational modifications (Liu *et al.*, 2008). In light of ARF biology such mechanisms might provide an additional layer of specificity determination in auxin output control. An example of ID in contribution to signaling diversity is the p53 tumor suppressor, which is involved in a wide set of cell fate decisions. Both the N- and C-terminal domains (comprising a third of the total protein sequence) are intrinsically disordered and contribute to most of the know protein-protein interactions (Dunker *et al.*, 2008). Furthermore, most of the post-translational modifications cluster on the intrinsic disordered regions (Dunker *et al.*, 2008). Besides a role in signaling diversity, intrinsically disordered domains can affect DNA binding. For example, the Drosophila transcription factor Ultrabithorax (Ubx) contains two intrinsically disordered domains that modulate the binding affinity of the structured DNA binding homeodomain (Liu *et al.*, 2008; Hsiao *et al.*, 2014).

The steroid hormone receptor (SHR) family is another class of proteins exemplifying the importance of ID in signaling. Similar to the MR of ARFs, the N-terminal transactivation domain (NTD), which can either activate or repress transcription, shows the least sequence homology among the SHR family and no structure of this region is available (Gallastegui *et al.*, 2015). The SHR have a modular structure and among 400 analyzed vertebrate and invertebrate SHR family members the NTD showed the highest level of disorder (69%) (Krasowski *et al.*, 2008). Induced folding of the NTD upon co-factor binding has been shown for the androgen-receptor (Reid *et al.*, 2002; McEwan *et al.*, 2007; Tantos *et al.*, 2012). Similar to p53, most post translational modifications fall within the NTD of SHR proteins (Lavery and Mcewan, 2005; McEwan *et al.*, 2007). The nature and convergence of different types of regulation on the ID domains implicate a focal point of extensive signal enhancement/diversity. To elaborate on the presence of intrinsic disorder, ARF protein sequences were analyzed using the disordered prediction algorithm PONDR-FIT (Xue *et al.*, 2010). The prediction, quite strikingly, shows a high degree of disorder in the MR of class A ARFs, which also seems to be conserved in the liverwort *Marchantia polymorpha* (Figure 2). There is a strong contrast to class B/C ARFs, which do not show this strong predicted disorder. Although there is no functional data supporting the existence of intrinsic disorder in the MR of activator ARFs, it provides a new concept in the explanation to the wide set of responses an ARF can elicit in specific cell types in response to auxin. Functional analysis of these ID regions should also help to define if ID is connected to the ability to activate gene expression.

**Figure 2:**
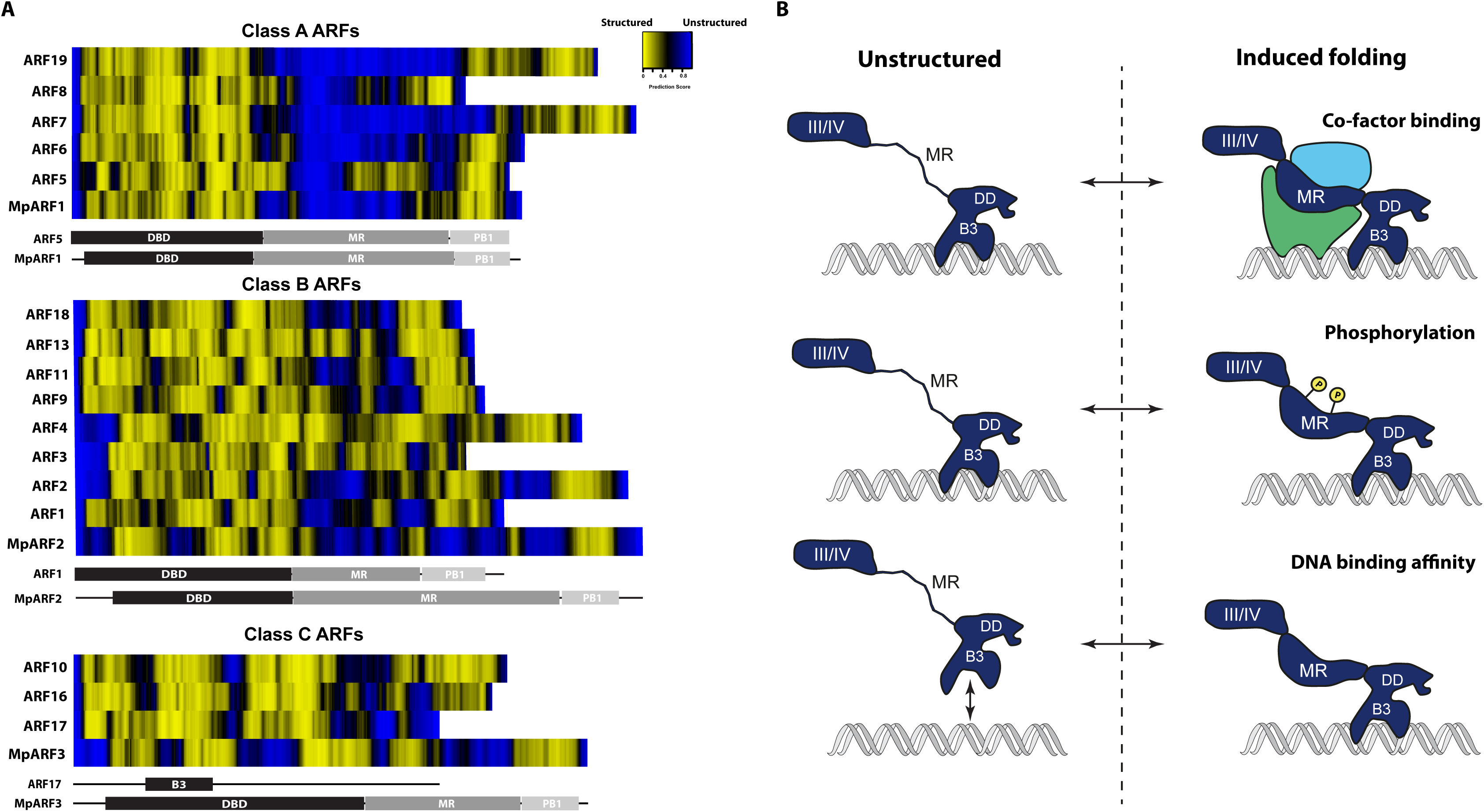
Intrinsic disorder in the ARF middle region. (A) Predicted disorder in the middle region appears to be a prominent and conserved feature in the class A “activator” ARFs. Full-length protein Arabidopsis ARF sequences, as well those from *Marchantia polymorpha* (MpARF) were used as input in the disorder prediction tool DisProt using the PONDR-FIT algorithm (Xue *et al.*, 2010). Disordered values were used in R to generate a heatmap. Domain locations were retrieved from UniProt. (B) Disordered regions can serve as a focal signaling hub by obtaining induced structure with cofactors, modulation by posttranslational modifications or aid in DNA binding affinity/specificity. *Protein abbreviations: ARF, AUXIN RESPONSE FACTOR; III/IV;C-TERMINAL PHOX AND BEM 1 DOMAIN; MR, MIDDLE REGION, DD, DIMERIZATION DOMAIN.*

### Regulation of ARF activity through the C-terminal domain

It has long been known that the C-terminal ARF domain mediates interactions with Aux/IAA proteins (Ulmasov *et al.*, 1997b). Structural analysis on the C-terminal domain recently revealed the structural basis of such heterotypic interaction of ARF5 (Nanao *et al.*, 2014), ARF7 (Korasick *et al.*, 2014), IAA17 (Han *et al.*, 2014) and PsIAA4 (Dinesh *et al.*, 2015). The structural analysis of ARF5 and ARF7 revealed type I/II PB1 domains and the chemical basis of dimerization (Korasick *et al.*, 2014; Nanao *et al.*, 2014). The domain has both acidic and basic motifs, which form a tertiary *β*-grasp-fold structure. The sidedness of the structure, with an acidic and a basic face that can interact with other PB1 domains via electrostatic interactions, creates a front to back arrangement. This arrangement underlies homo- and hetero-dimerization between ARFs and with Aux/IAAs that also carry a PB1 domain and use it to interact with ARFs.

Several studies explored interaction specificity between Aux/IAA and ARF proteins, in an effort to map pathway complexity that might explain diverse auxin outputs. Two comprehensive studies utilizing large scale yeast 2-hybrid (Y2H) assays showed the variety at which these interactions can occur (Vernoux *et al.*, 2011; Piya *et al.*, 2014). Interestingly, in this assay, class B and C ARFs have limited to no interactions with Aux/IAAs (Vernoux *et al.*, 2011; Piya *et al.*, 2014). This suggests that auxin regulation within the nuclear pathway exclusively converges upon class A ARFs. Taken at face value, this finding would suggest that class B and C ARFs are disconnected from auxin regulation, and act by counteracting class A ARFs, for example by competing for DNA binding or blocking through heterodimerization (Richter *et al.*, 2013). It should be noted that in these large-scale interaction studies, proteins are expressed at much higher levels than naturally occurring and might also have increased stability. From studies in the moss *Physcomitrella patens,* a model was suggested wherein class A and B ARFs either compete or cooperate to repress or induce transcription respectively (Lavy *et al.*, 2016). It appears that more *in vivo* studies are dearly needed to determine if and how class B and C ARFs are wired into the auxin response network, and what purpose their PB1 domains have.

An interesting finding in the structural analysis of ARFs and Aux/IAA proteins was that PB1 domains have the capacity to oligomerize *in vitro,* in crystal and in solution (Korasick *et al.*, 2014; Nanao *et al.*, 2014). The biological significance of such oligomerization is still an open question. ARF5 that lacks the PB1 domain has reduced capacity to bind DNA *in vitro,* and this could be overcome by antibody-induced dimerization (Ulmasov *et al.*, 1999a). Thus, PB1-interactions, in addition to being the site for auxin regulation through Aux/IAA binding, could potentiate DNA binding. Mathematical modeling of TIR1/AFB, auxin, ARF and Aux/IAA interactions provide a conceptual basis for significance of ARF oligomerization on auxin output (Farcot *et al.*, 2015). Aux/IAA-ARF interactions may determine the amplitude, Aux/IAA-Aux/IAA interactions the speed and ARF-ARF interactions the sensitivity of the response. Since the parameters depend on the PB1 domain interaction, oligomerization may significantly affect the auxin output (Weijers and Wagner, 2016). On the other hand, questions can be raised about the relevance of mediated ARF DNA binding by the homo/hetereodimerization through the PB1 domain. For example the truncated ARF5 (ΔPB1) is hyperactive and still able to activate transcription (Krogan *et al.*, 2012). Also, ARF4 and ARF3 act redundantly in establishing leaf polarity (Pekker *et al.*, 2005). Since ARF3 naturally lacks a PB1 domain it appears that this domain is not required for ARF function in this context. A kinetic analysis of ARF-ARF, ARF-Aux/IAA and Aux/IAA-Aux/IAA interactions *in vitro* showed that the affinity of ARF:ARF homo-dimers is ∼10 to ∼100 fold lower than ARF:AuxIAAs hetero-dimers (Han *et al.*, 2014). This suggests that equilibria will tend to favor heterotypic interactions, thus endowing auxin regulation upon ARFs.

## Dynamic control of auxin-dependent genes in a chromatin context

An important question is how auxin – and ARFs – can regulate genes in the context of chromatin. It had previously been shown that Aux/IAA proteins recruit the co-repressor TOPLESS (TPL), and likely repress expression through histone de-acetylation (Long *et al.*, 2006; Szemenyei *et al.*, 2008). Recently, a chromatin switch mechanism has also been proposed to direct ARF-dependent gene activation. Chromatin can be configured in a bipartite manner; either closed marking an inactive state or an open configuration marking an active state. Recently a switch in this state was found in which ARF5 is able to unlock closed chromatin in concert with the SWI/SNF chromatin remodelers BRHAMA (BRM) and SPLAYED (SYD) (Wu *et al.*, 2015). Aux/IAA proteins compete with SWI-SNF recruitment to ARF5, and thus Aux/IAA degradation allows chromatin remodeling (Wu *et al.*, 2015). Furthermore, the GRE motif-binding bZIP transcription factors can recruit the histone acetyltransferase (HAT) SAGA complex to a GH3 gene and induce auxin responsive transcription (Weiste and Dröge-Laser, 2014). Interestingly, a conserved bZIP motif was shown to be occluded prior to ARF5-dependent chromatin unlocking (Wu *et al.*, 2015). From these two studies it follows that there may be a concerted action of ARF5-induced nucleosome remodeling followed by HAT-dependent histone modification during developmental reprogramming. Since this mechanism has so far only been demonstrated for ARF5, it will be interesting to see if all class A ARFs, and possibly class B/C ARFs, operate in a similar manner.

Conversely, it was recently shown that histone deacetylation plays a role in the regulation of genes by other class A ARFs (Fukaki *et al.*, 2006). The ARF7/19 and IAA14 proteins play a critical role in lateral root initiation (Okushima *et al.*, 2005). Through phenotypic analysis and exogenous histone deacetylase inhibitor application it was shown that the chromatin remodeler PICKLE (PKL) and histone deacetylation are required for IAA14-mediated ARF7/19 inhibition. Since PKL strongly resembles the mammalian CHD3/Mi-2 protein of the Nucleosome Remodeling Deacetylase complex (NuRD), consisting of several histone deacetylases, it is conceivable that such concerted action of remodeling and histone deacetylation takes place on ARF target loci.

Interactions between ARFs and chromatin regulators appear to be multi-layered and complex. For example, under low auxin levels, the TPL co-repressor bridges the CDK8 kinase module (CKM) of the MEDIATOR complex with the ARF7/19 - IAA14 module (Ito *et al.*, 2016). The CKM Mediator module prevents the association of the core Mediator subcomplex with RNA polymerase II (Ito *et al.*, 2016). The TPL-mediated interaction is probably distinct from the proposed recruitment of histone de-acetylases by TPL (Long *et al.*, 2006), and importantly it might not involve covalent histone modifications. Under high auxin levels, IAA14 becomes degraded thus leading to loss of the TPL-CKM bridge followed by active transcription (Ito *et al.*, 2016). Such a sequences of events resembles a primed transcriptional state that can accommodate quick transcriptional responses. It is clear from the few examples given here that we are only beginning to scratch the surface of chromatin-level control in ARF action, and further exploration in this area is likely to give much more insight into the fast and dynamic regulation of auxin-responsive genes.

## No protein is an island – ARF cofactors shape auxin response

Other than interaction with chromatin regulators, transcription factors (TF) usually cooperate with co-factors that can modulate DNA binding specificity or transcriptional activity. Such interactions can assemble into higher-order protein complexes that can regulate the local chromatin environment and activate or repress gene transcription. In some instances, as reported for the Drosophila Hox TFs, co-factors can modulate the TF to gain novel DNA binding specificities (Slattery *et al.*, 2011). In comparison with other TFs, the number of reported co-factors for ARFs is limited and, if reported, the precise functionality of the interaction not completely elucidated (Figure 3). Since co-factors are important in modulating TF activity, it is conceivable that ARF co-factors play a significant role in modulating activity.

**Figure 3:**
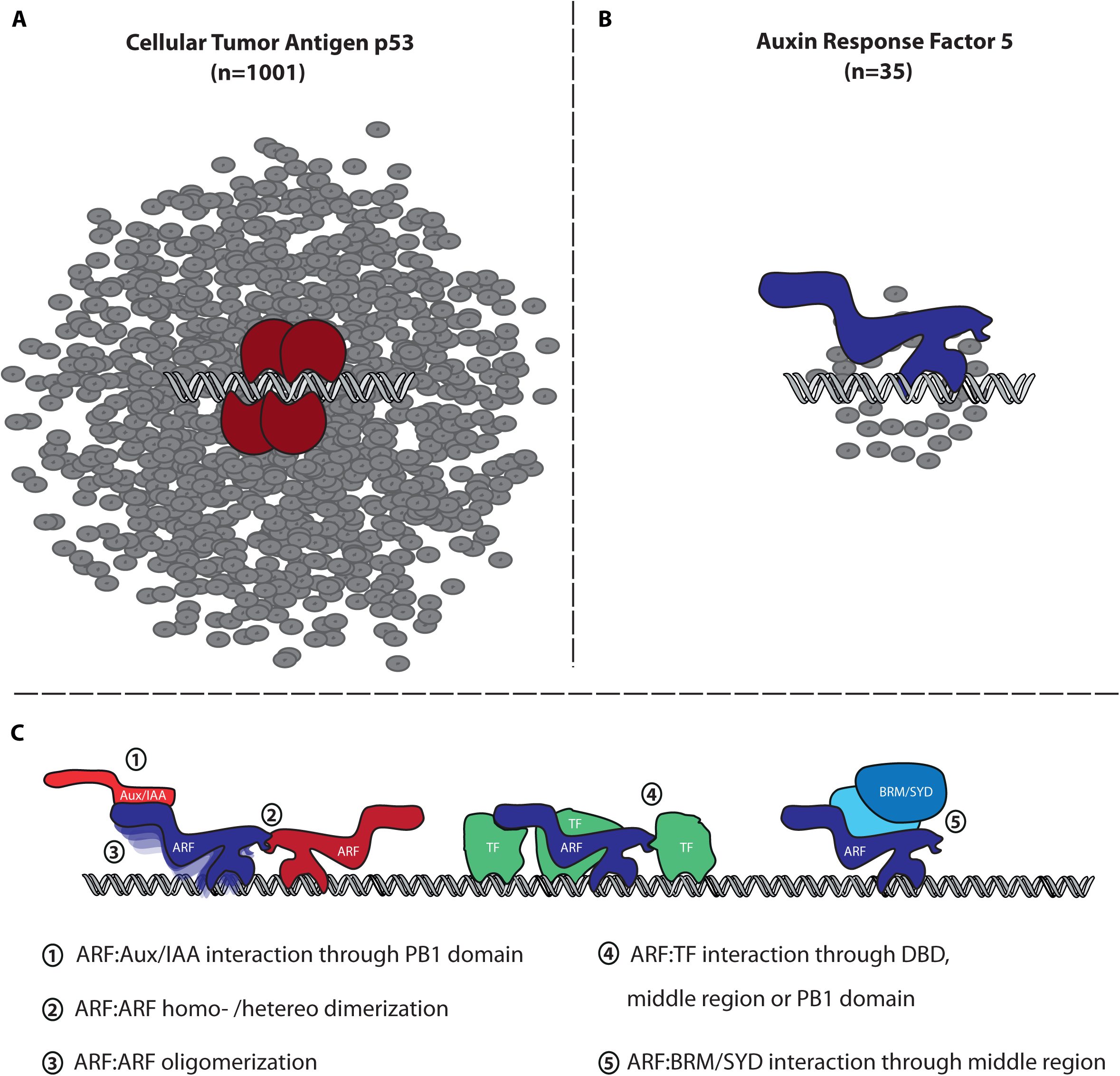
ARF cofactors. (A,B) Complete interactome of the human tumour suppressor p53 (A) and ARF5 (B) depicts the limited state of our knowledge on ARF functioning in comparison with p53. Figure was made utilizing Cytoscape by selecting direct neighbours and using the BioGrid database (last accessed march 2017). (C) Current known modes of interactions and interactions surfaces of ARFs. *Protein abbreviations: ARF, AUXIN RESPONSE FACTOR; Aux/IAA, AUXIN/INDOLE-3-ACETIC ACID, BRM, BRAHMA; SYD, SPLAYED; TF, TRANSCRIPTION FACTOR*

Interactions between TFs can occur within and between families (Bemer *et al.*, 2016). For ARFs, such (ARF-ARF) interactions have only been shown *in vitro* and appear to be a requirement for high-affinity DNA binding (Boer *et al.*, 2014). Interactions between transcription factors of different families are also frequently reported, extending the repertoire of TF activity and integrating several developmental, environmental and hormonal pathways. For ARFs this has been shown in several instances. An example is the interaction between MYB77 and ARF7. It was shown that this interaction is important for the regulation of auxin-dependent genes and might integrate abscisic acid signaling with auxin response (Shin *et al.*, 2007; Zhao *et al.*, 2014). A more complex integration was shown for ARF6, which interacts with the bHLH factor phytochrome interacting factor 4 (PIF4) and brassinazole resistant 1 (BRZ1) to regulate a common set of target genes (Oh *et al.*, 2014). It was further shown by genetic studies and Y2H that gibberellic acid signaling integrates in the ARF6-PIF4-BZR1 complex by disrupting ARF6-PIF4 interaction through the DELLA protein repressor of GA (RGA). Of note is that the PIF4 and RGA interactions predominantly occur through the middle region and that RGA also interacts with ARF7 and ARF8 (Oh *et al.*, 2014). Another bHLH (big petal (BPE)) - has also been shown to support ARF function. ARF8 and BPE synergistically act during petal organ growth (Varaud *et al.*, 2011). It was further shown that ARF8, but also ARF6, interacts with the MADS-box transcription factor FRUITFULL (FUL) to promote fruit valve growth (Ripoll *et al.*, 2015). Although the primary focus of the described ARF-TF interactions all relate to class A ARFs interactions with class B ARFs have also been described to a lesser extent. For example, ARF3 has been studied in the context of polarity determination where it interacts with the GARP family member ABERRANT TESTA SHAPE. In two studies, ARF2 has been shown to interact with MADS-box TF FUL and AP1 (Smaczniak *et al.*, 2012; Ripoll *et al.*, 2015).

From this non-exhaustive list of examples, it is apparent that ARFs are not the sole entities in regulating auxin dependent transcription. One prominent question that can be raised from the studies reported thus far is whether there is a common mode of regulation on auxin target genes. It appears that hetereotypic TF interactions are common, especially for class A ARFs. Cooperative DNA binding of two TFs can result in a net increase in affinity for their motifs while the specificity for the motifs remains unchanged (Spitz and Furlong, 2012). On the other hand cooperative binding can also create new specificities. It appears that cooperative binding plays a role in ARF dependent transcriptional activity as is the case for many other plant related TFs (Bemer *et al.*, 2016). MYB77 has interaction with ARF7 and bZIP-dependent SAGA complex recruitment induces auxin transcription (Shin *et al.*, 2007; Weiste and Dröge-Laser, 2014). The binding motifs of MYB and bZIP have been shown to be enriched and evolutionary conserved near AuxRE (Berendzen *et al.*, 2012).

Currently a comprehensive analysis on ARF/cofactor interactions is lacking. An unbiased *in planta* approach on all ARFs, as was for example performed on several MADS-box TFs (Smaczniak *et al.*, 2012), could promote our understanding on how ARFs regulate transcription. In perspective, the BioGrid interaction database lists over 1000 interactions for the human p53 protein while ARFs only have a small portion of that number listed (Figure 3). This exemplifies that the field is currently far from understanding ARF biology.

## Is it really that simple?

Historically, ARF1 was first found in a yeast 1-hybrid screen to identify transcription factors which bind on a synthetic DNA (P3[4x]) known to be highly auxin-responsive (Ulmasov *et al.*, 1997a). All others ARFs have been found by sequence homology to ARF1 (Guilfoyle *et al.*, 1998). This history urges an existential question: are all ARFs really ARFs? Do all ARFs mediate auxin response? Is an ARF that is not able to interact with Aux/IAA proteins still connected to the auxin response network? The PB1 domain is lacking in ARF3, ARF13, ARF17, and ARF23. ARF23 is different from all others as it is heavily truncated from its DBD. It has been show that deletion between DBD and MR can affect dimerization of ARF18 (Liu *et al.*, 2015), so there is good chance that ARF23 is not able to dimerize. Moreover its biological function or its ability to bind DNA is not known, and given that this gene is part of a recently duplicated cluster near the centromere of chromosome I (Okushima *et al.*, 2005), there is a chance that ARF23 is becoming a pseudogene.

For ARF3 and ARF17, it appears that despite lack of the PB1 domain, these proteins do control auxin-dependent development (Mallory *et al.*, 2005; Simonini *et al.*, 2016). Y2H showed that ARF17 was able to interact with Aux/IAAs, despite it is lacking the conserved PB1 (Piya *et al.*, 2014). Moreover, truncated ARF5 or ARF7 (lacking the PB1 domain) could still be activated by auxin, though less efficiently than the full-length protein (Wang *et al.*, 2013). Even if *in planta* proof is lacking, these findings raise the possibility that Aux/IAAs can even interact with truncated ARFs. Thus, it appears that the lack of PB1 can not be used as a criteria to discriminate ARF from non-ARF.

In the past decades, research efforts characterized the canonical auxin signaling pathway wherein, under high auxin levels, repressive Aux/IAAs become degraded, relieving ARFs from repression. Although this auxin perception mechanism is well known, the regulatory mechanism by which ARFs control auxin output is still vaguely understood. Another aspect that is not currently investigated is the biological relevance of ARF heterodimerization. Few studies have demonstrated the ability of distinct ARFs to interact *in vitro*. Heterodimerization has been observed in gel shift assays between ARF1 and ARF4 (Ulmasov et al. 1999) or between different ARFs in Y2H experiments (Ouellet *et al.*, 2001; Hardtke *et al.*, 2004; Vernoux *et al.*, 2011). While it is thus clear that ARFs can heterodimerize, it needs to be established whether they do so *in vivo,* and the biological relevance of heterodimerization must be understood.

Besides the mechanism that concern the homeostasis of the nuclear auxin pathway, recent research revealed non-canonical pathways that effect ARF regulated gene expression. In the canonical pathway, control by posttranslational modifications have been identified, such as *cis-trans* proline isomerization of Aux/IAAs (Dharmasiri *et al.*, 2003), S-nitrosylation of TIR1 (Terrile *et al.*, 2012) and phosphorylation of Aux/IAAs (Colón-Carmona *et al.*, 2000). For ARFs, phosphorylation events have been shown to be important for their function. During low potassium availability the K+ transporter HAK5 is upregulated to compensate for K+ deficiency (Gierth *et al.*, 2005). The control of the HAK5 gene is modulated by ARF2. In the presence of sufficient K+ levels, ARF2 represses HAK5 transcription (Zhao *et al.*, 2016). In K+ deficiency environments ARF2 becomes phosphorylated blocking ARF2 DNA binding activity (Zhao *et al.*, 2016). This mechanism of modulation of DNA binding activity by phosphorylation has been shown on ARF2 by the brassinosteroid (BR) -regulated BIN2 kinase (Vert *et al.*, 2008). The integration of BR signaling components and activity modulation on activator ARFs has also been reported (Cho *et al.*, 2014). During lateral root organogenesis ARF7 and ARF19 play pivotal roles and it was shown that the auxin module does not solely control the activity of these ARFs during this process. The BIN2 kinase phosphorylates these ARFs and inhibits Aux/IAA interaction potentiating ARF activity (Cho *et al.*, 2014). Quite surprisingly is that BIN2 in this process is not activated by BR but by the tracheary element differentiation inhibitory factor (TDIF) peptide (Cho *et al.*, 2014).

Other than phosphorylation, a recent finding revealed an alternative auxin sensing mechanism resembling the animal thyroid hormone receptor pathway. The atypical (class B) ARF3/ETT is involved in auxin regulated gynoecium patterning (Sessions *et al.*, 1997; Simonini *et al.*, 2016). Since ETT lacks a PB1 domain, canonical auxin signaling is not likely to regulate ETT activity. ETT interacts with the basic helix-loop-helix (bHLH) transcription factor INDEHISCENT (IND) and this interaction is auxin-sensitive (Simonini *et al.*, 2016). In a bimolecular fluorescence complementation experiment, upon addition of auxin, the ETT:IND dimer appeared to dissociate. Further Y2H experiments showed similar results for the ETT:IND dimer but also for other ETT:TF dimer complexes (Simonini *et al.*, 2016).

These results show how elaborate ARF activity can be modulated beside the core nuclear auxin module. An interesting question is whether these non-canonical pathways represent a general mode of action in ARF activity modulation.

## Concluding remarks

The past few years, many studies gave new details about ARFs mode of action and functions of their conserved domains. They confirmed the key role of the ARF as an output of the nuclear auxin pathway but particularly emphasizes new characteristics of ARF that were not suspected before. The mode of action of the ARFs was seen more like an on/off mechanism on TGTCTC motif while now, it is believed that ARF are more flexible than that and could be part of larger protein complex (chromatin switch or TF-TF). However, these recent breakthroughs raise new questions and need to be challenged first. Even if these findings brought new insights into ARF mode of action, it is still difficult to give a precise definition to describe this family. One of the reasons is that only little is known about the universality of these mechanisms. Testing these hypothesis on different ARFs classes (A,B,C) or “activators”/”repressors” ARFs will probably help to draw a mugshot of an ARF. It is also worth to highlight that some ARFs still have not been biologically characterized. It will be necessary to extend this knowledge to other species phylogenetically distant from *Arabidopsis* in order to understand how the auxin signaling pathway has evolved into a complex and apparently fine tuned system.

## Acknowledgements

We apologize to colleagues whose important contributions we could not discuss in this review. Research on auxin response in D.W.’s lab is funded by the Netherlands Organization for Scientific Research (NWO; VICI 865.14.001). S.P. is supported by a long-term fellowship from the European Molecular Biology Organization (EMBO; ALTF-535-2014) and the Bettencourt Schueller Foundation.

